# A Comprehensive Atlas of Cell Type Density Patterns and Their Role in Brain Organization

**DOI:** 10.1101/2024.10.02.615922

**Authors:** Rodrigo Muñoz-Castañeda, Ramesh Palaniswamy, Jason Palmer, Rhonda Drewes, Corey Elowsky, Karla E. Hirokawa, Nicholas Cain, Kannan Umadevi Venkataraju, Hong-Wei Dong, Julie A. Harris, Zhuhao Wu, Pavel Osten

## Abstract

Cell-type composition across brain regions is a critical structural factor shaping both local and long-range brain circuits. Here, we employed single-cell resolution imaging of the mouse brain, combined with computational analyses, to map the distribution of 30 cell classes and types defined by gene marker expression in Cre recombinase-based genetic mouse models. This approach generated a comprehensive atlas of cell type-specific densities across the male and female brain, revealing (1) surprisingly broad sex differences in cells tagged by developmental cell-type markers, (2) shared cell type composition signatures among functionally related brain structures, and (3) close associations not only between specific cell types but also discrete cell type densities and anatomical regions and subregions. In summary, despite the relatively broad cell type classification enabled by the Cre mouse models, our findings highlight intricate relationships between brain cell type distribution and anatomical organization, associating distinct local cell densities with region-specific brain functions.

## INTRODUCTION

The mammalian brain consists of numerous distinct cell types, each distributed in specific densities and organized into cell type-specific circuits within anatomically and functionally defined regions. Since cell type composition establishes the structural foundation for the formation of function-specific neural circuits, it plays a pivotal role in supporting all brain functions, from information processing and integration to the regulation of behavior and emotional responses^1,2^.

The classical anatomical division of the mammalian brain includes seven major structures: the cortex, cerebral nuclei, thalamus, hypothalamus, midbrain, hindbrain, and cerebellum. The spatial organization of the subregions within these structures has been characterized in brain atlases using histological techniques such as Nissl staining, which visualizes brain regions based on differences in cell density and size, or acetylcholinesterase (AChE) immunolabeling, which highlights structures based on cholinergic innervation. While cytological staining, immunolabeling, and anterograde and retrograde projection tracing have been invaluable for mapping brain regions, traditional anatomical approaches are limited by their reliance on a small number of brain samples, failing to account for inter-subject variability, and the potential for expert bias. Consequently, discrepancies exist among anatomical atlases, particularly in mouse brain atlases based on Nissl and AChE staining, Nissl alone, and more recent methods employing highly averaged brain autofluorescence^3–5^.

Efforts to reconcile the anatomical atlas differences have included a consortium-based study integrating multimodal data from various laboratories to provide detailed descriptions of specific anatomical areas, such as the mouse motor cortex^6,7^, as well as recent digital integration of multiple atlases within the Common Coordinate Framework (CCF) of the mouse brain^4^. Given the importance of the mouse as a model organism in basic neuroscience—particularly its utility for tracing and manipulating specific cell types using cell type-specific Cre recombinase mouse lines^3,8^—it remains important to further examine the relationship between mouse brain anatomical organization and its structural organization defined by the distribution of different cell types.

In this study, we utilized 30 cell type-specific Cre mouse lines (see Table 1) to quantitatively map the respective cell type populations in the adult brain, providing a comprehensive data resource for conducting anatomical and functional studies using these mouse lines. The Cre mouse lines include twenty-seven widely used cell type-specific knock-in Cre lines, where Cre recombinase is expressed from an IRES cassette inserted after the stop codon in the 3′ region of the cell type marker gene^9^. This design achieves faithful Cre expression alongside the native gene product. Additionally, we included three commonly used bacterial artificial chromosome (BAC) transgenic mouse lines where Cre expression from the Ntsr1 (neurotensin receptor 1), Rbp4 (retinol-binding protein 4), and Tlx3 (T-cell leukemia homeobox 3) promoters highlights different cortical layers^10,11^.

Using these data, we first show that while the total neuronal numbers do not differ between male and female brains, sex-dimorphic differences exist in the numbers of neuronal populations labeled by developmentally expressed genes, such as the transcription factors Rbp4 (Retinol Binding Protein 4), Cux2 (Cut-Like Homeobox 2), Rorb (RAR-Related Orphan Receptor Beta), Tlx3 (T-Cell Leukemia Homeobox 3), and the vesicular glutamate transporters VGlut2 and VGlut3, suggesting gene- and cell population-specific sex-based differences in brain development. Next, we applied unbiased clustering of major brain regions based on the similarity of their cell type composition to reveal that both gross anatomical areas and cortical and other subregions can be defined by shared composition of specific cell types, linking cell type structural signatures to distinct brain functions. Finally, we developed a novel approach to analyzing whole-brain cell type distributions using Density-Based Spatial Clustering of Applications with Noise (DBSCAN), an algorithm that partitions spatial data based on local densities. This unbiased approach revealed that cell type-based cell densities can define anatomical borders, link brain subregions with shared functions, and identify putative novel subareas with distinct roles. For example, distinct cell densities of cortical pyramidal neuron markers VGlut1, Rbp4 and Cux2 delineate frontomedial, motor, lateral association and sensory cortical areas. Similarly, the dentate gyrus (DG), CA1, CA2, CA3 as well as the fasciola cinereum (FC) of the hippocampal formation can be spatially identified based on VGlut1, VGlut2, Calb2 (calretinin) and SERT (serotonergic transporter) cell densities. Applying the same cell density-based parcellation across the brain using diverse cell type markers revealed input-matching parcellation of the dorsal striatum as well as nuclei-specific parcellation of the thalamus and hypothalamus.

## RESULTS

### Whole brain visualization and quantification of gene expression-defined brain cell types

The whole brain mapping approach applied in the current study included the following: First, we crossed genetically engineered “Cre-driver” mouse lines expressing the Cre recombinase from cell type-specific promoters with “Cre-reporter” mice expressing nuclear H2B-GFP protein (or tdTomato protein) upon Cre-mediated recombination. This resulted in nuclear fluorescent labeling of the gene expression-defined cell types in all 30 double-transgenic Cre-lox mice (Figure 1a). The whole mouse brain imaging was done by serial two-photon tomography (STPT) and the resulting serial section datasets were registered and computationally analyzed using our whole-brain mapping platform^3,6,12^, with new modifications to enhance the precision of data registration, support data visualization at a full image resolution and enable density clustering by DBSCAN for of each cell type (Figure 1b-g, Methods). The resulting 280-coronal section datasets (XY resolution 1 μm, Z-spacing of 50 μm) were registered onto the mouse brain Common Coordinate Framework (CCF)^3,5^, and the fluorescent nuclei were detected by convolutional neural networks (CNNs), and all processed datasets were visualized in 3D point-cloud views (Methods). This revealed distinct anatomical patterns for each Cre-based cell type, such as broad and fairly uniform patterns for neuron-specific SNAP25 or glia cell-specific Olig2, broad but varied patterns for inhibitory neuron-specific GAD2 or Slc32, inhibitory subtype-specific PV, Calb, Calb2, SST, VIP and Pdyn, and excitatory neuron-specific VGluT1, VGluT2, VGluT3 and Emx1, as well as more distinct patterns of the Avptm gene marking hypothalamic vasopressinergic cells, the Oxt marking oxytocinergic cells, or ChAT marking cholinergic neurons (Figure 2a and Supplementary Videos 1-4). All datasets are deposited at the Brain Image Library for public sharing https://www.brainimagelibrary.org/).

**Figure 1:**
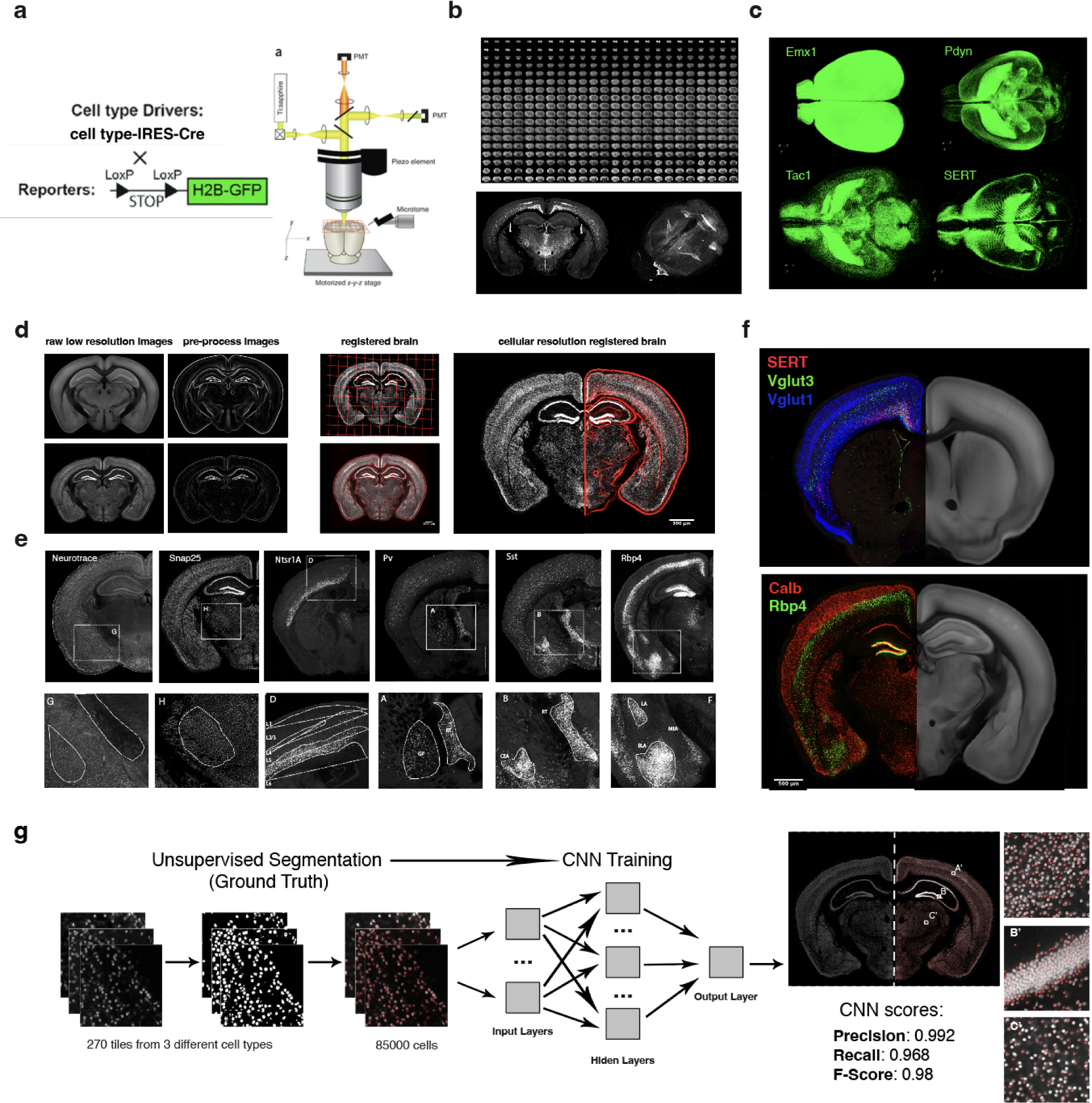
**Whole brain analysis and visualization at cellular resolution**: **a)** Serial Two Photon Tomography (STPT) imaging of mouse brains from cell type Cre-driver lines crossed with fluorescent protein Cre-reporter lines. (Left) A diagram of Cre-LoxP labeling strategies. (Right) A schema of STPT instrument for automated whole mouse brain imaging. **b)** (Top) Resulting 280 serial sections of whole mouse brain imaging at 1x1x50 um resolution. An example coronal section and the 3D renderization of the whole brain. **c)** Examples of 3D renderization of fluorescent H2BGFP cell distribution for Emx1, Pdyn, Tac1 and SERT Cre-driver lines. **d)** Intrinsic anatomical features (IAFs) based 3D registration of mouse brain at cellular resolution. (Left) Generation of anatomical features from downsampled 20x20x50 um whole brain image for both mean intensity reference brain(top) and an individual sample brain. (Middle) The resulting grid space deformation after registration (top) and the overlay of the STPT imaged brain onto the reference IAFs based brain (bottom). (Right) The resulting transformations obtained from the downsampled images were applied to the original 1x1x50 um resolution brain images and warped onto a high-resolution reference space coordinate system. **e)** Examples of NeuroTrace and different Cre-driver lines crossed with the H2BGFP Cre report showing cell type-specific anatomical distribution before registration. **f)** Overlay of selected 4 Cre-driver lines SERT+VGlut3+VGlut1 (top) and Calb2+Rbp4 (bottom) after registration at the 1x1x50 um cellular resolution (left) onto the reference brain (right), showing their cellular distribution organization. **g)** (Left panel) Machine learning-based cell detection. Unsupervised segmentation strategy was used for the generation of a large dataset of marked cell nuclei from 3 different imaged brains used for training of a convolutional neural network (CNN). Resulting accuracy was validated using manually marked up ground truth data. (Right panel) Coronal plane of Snap- H2B-GFP with left hemisphere overlay onto the output segmentation, with details showing the segmentation results at the cortex (A’), hippocampal formation (B’) and the thalamus (C’).

**Figure 2:**
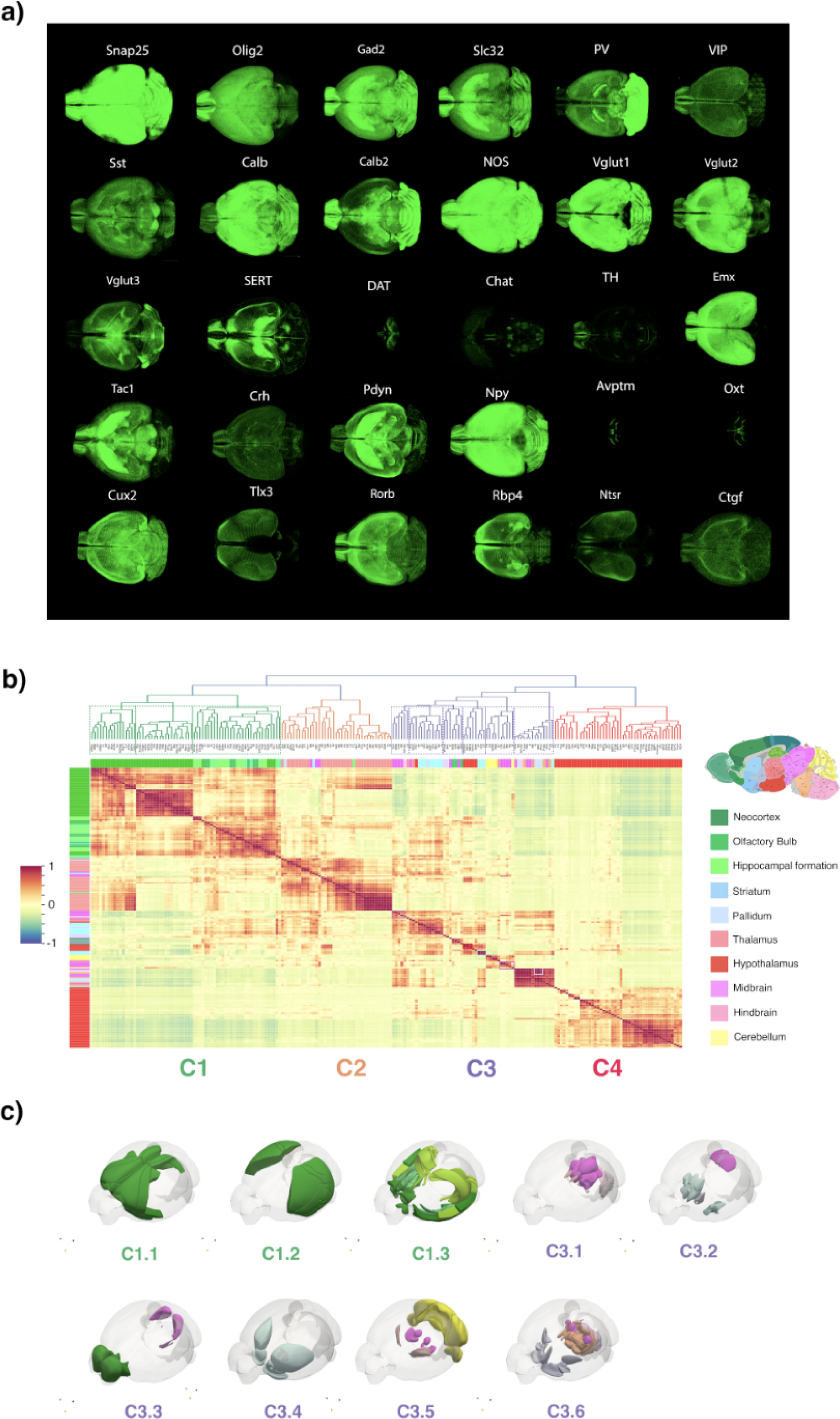
**Whole brain 3D distributions visualization of cell types and analysis**: **a)** 3D point-cloud visualizations from 30 different cell types: Snap25 - Synaptosomal associated protein 25-; Olig2 - Oligodendrocyte Transcription Factor 2-; Gad2 - Glutamate Decarboxylase 2-; Slc32 - Vesicular GABA transporter-; Pv -Parvaobumin-; VIP - Vasoactive Intestinal Peptide-; Sst - Somatostatin-; Calb -Calbindin-; Calb2 -Calretinin-; NOS - Neuronal nitric oxide synthase 1-; VGlut1 - Vesicular glutamate transporter 1-; VGlu2 - Vesicular glutamate transporter 2-; VGlut3 - Vesicular glutamate transporter 3-; SERT -Serotonin Transporter-; Chat - Choline acetyltransferase -; DAT - Dopamine Transporter -; TH - Tyrosine Hydroxylase-; Emx1 - Empty Spiracles Homeobox 1-; Tac1 - Tachykinin precursor 1-; Crh - Corticotropin-releasing hormone-; Pdyn - Prodynorphin-; Npy -Neuropeptide Y-; Avptm - arginine vasopressin-;-; Oxt - Oxytocine-; Cux2 - Cut Like Homeobox 2-; Tlx3 - T Cell Leukemia Homeobox 3; Rorb - RAR Related Orphan Receptor B-; Rbp4 - Retinol binding protein 4-; Ntsr1A - Neurotensin receptor type 1-; and Ctgf - Connective tissue growth factor-. **b)** Clustermap showing mouse brain organization based on the distribution of all the cells. Cluster analysis shows that brain areas organize in 4 major clusters based on their cell distribution: cortex (cluster C1, green dendrogram), thalamus (cluster C2, orange dendrogram); striatum, pallidum, midbrain, hindbrain and cerebellum (cluster C3, purple dendrogram) and hypothalamus (cluster C4, red dendrogram). Thus, major brain has a very distinctive cellular organization defining their internal structure. **c)** Normalized cell type distribution on major brain areas. X-axis show all cell types clustered by normalized expression, and Y-axis shows major brain areas organized by panel B) brain cluster organization.

Since the Cre reporter marks cells expressing the Cre protein at any time during development till the adulthood^13^, we first determined which labeled cell-type distributions represent adult expression of the marker gene and which include additional development-based expression that is not maintained in the adult brain. This was done by comparisons of the cell type fluorescent nuclei distribution in our datasets to *in-situ* hybridization data from the Allen Institute for Brain Science data portal^14^, the Gene Expression Nervous System Atlas (GENSAT) portal^15^, and previously published cell type labeling studies^16,17^ (Supplemental Figure 1). This revealed that the Cre-based nuclear labeling largely agrees with adult gene expression in 26 Cre-lox mouse lines, while 4 lines—the SERT, VGlut2, VGlut3 and nNOS (neuronal nitric oxide synthase)—comprise strong developmental labeling in addition to the adult expression: 1) native nNOS expression marks a sparse population of inhibitory neurons in the adult cortex and other brain areas, but nNOS-Cre-based labeling marks high density of cells across the entire brain, reflecting a strong nNOS developmental expression^18^; 2) native SERT expression marks serotoninergic neurons in the adult midbrain nuclei^19,20^, but the SERT Cre-based labeling marks dense neuronal populations specifically in cortex and the thalamus, reflecting SERT developmental expression during the formation of cortico-thalamic connection patterns^21,22^; 3) native VGlut2 expression marks adult glutamatergic neurons in subcortical areas^23,24^, but VGlut2-cre labeling also includes cortical areas, reflecting proposed VGlut2 role in the refinement of cortical pyramidal neuron dendritic arbors^24–26;^ and 4) native VGlut3 expression marks adult cholinergic neurons of the striatum^27^, a subpopulation of serotoninergic neurons in the dorsal raphe^28^, and at low density other neurons across all major areas^29,30^, but VGlut3-Cre labeling marks additional distinct cell populations in the cortex and the thalamus^31^ (see Supplementary Information and Discussion).

The full quantification of Cre-based cell types across anatomical brain regions of the Allen Mouse Brain Atlas (ARA) is provided as a resource for the neuroscience community in Table 2. This quantification also enabled us to statistically compare cell type distribution between male and female brains, revealing significant sex-based differences in several cell type-specific Cre mouse lines. While a more detailed analysis of these sex-based differences will be presented in a separate manuscript, we provide here, as a resource for the neuroscience community, all identified sex differences mapped within the ARA mouse brain atlas in Table 2, along with a brief description of our findings.

First, we note that we did not identify sex-based differences in SNAP25-positive cells, which represent the majority of brain neurons, or in VGluT1-positive cells, which represent the majority of cortical and hippocampal excitatory neurons. This suggests that there are no substantial differences in overall neuron counts between male and female brains, consistent with previous studies^32^. Second, in contrast to the SNAP25 and VGluT1 data, we identified prominent sex-based differences in specific cell types labeled by genes with known developmental expressions and functions. In particular, we found significantly more cells expressing the transcription factors (TFs) Cux2, Rbp4, Rorb, Tlx3, and Olig2 (Oligodendrocyte Transcription Factor 2), as well as the glutamatergic transporter VGluT2 in the male brain. Conversely, there were significantly more cells expressing the glutamatergic transporter VGluT3 in the female brain (Table 3). The transcription factors Cux2, Rbp4, Rorb, and Tlx3 all label excitatory neurons during cortical development. Specifically, Cux2 is most densely expressed in superficial layer 2/3 pyramidal neurons, Rbp4 and Tlx3 in layer 5 pyramidal neurons, and Rorb in both layer 4 and layer 5 pyramidal neurons. The expression of VGluT2 and VGluT3 in the cortex also reflects transient developmental functions, with VGluT2 marking broad populations of cortical and hippocampal pyramidal neurons, while VGluT3 is more restricted to specific cortical neuron populations. Taken together, these findings suggest that sex-specific developmental differences in gene expression, rather than differences in the total pyramidal neuron population, may contribute to sex-specific modulation of brain circuits and functions during development.

Finally, we observed significantly more cells expressing the oligodendrocyte marker Olig2 (oligodendrocyte transcription factor 2) across most brain regions in the male brain. While these differences also start occurring during the development, they appear to represent true increases in the number of oligodendrocytes across the entire male adult brain, supporting earlier results based on oligodendrocyte or myelin-related markers within selected brain regions, such as the corpus callosum, fornix, and spinal cord^33,34^.

### Shared cell type composition across gross anatomical brain areas

The distinct global cell distribution patterns revealed by the 3D data visualization in Figure 2a support the widely accepted, though not directly tested, concept that gross anatomical areas may be defined by their shared composition of specific cell types. To test this, we clustered major anatomical areas based on the similarity of their cell type composition, using a normalized expression of each cell type across brain areas (Figure 2b, see Methods). This analysis identified four major clusters: Cluster 1 comprised mainly cortical areas, Cluster 2 primarily included thalamic nuclei, and Cluster 4 consisted mainly of hypothalamic nuclei. In contrast, Cluster 3 encompassed a more diverse set of brain regions, including the striatum, midbrain, hindbrain, cerebellum, and olfactory areas (Figure 2b). Within the cortical cluster, three major subclusters were observed: (1) frontomedial association and motor areas, (2) lateral association and sensory areas, and (3) the cortical subplate, hippocampal formation, and olfactory areas (Figure 2b-c; see also Figure 3 for a more detailed cortical analysis). These findings demonstrate that the thalamus and hypothalamus each maintain a relatively homogeneous cell type composition across their respective subregions. Meanwhile, the cortical cell type composition further distinguishes brain areas involved in distinct functions, including executive control, decision-making, and emotion; sensory perception and multimodal integration; and memory processing, sensory development, and olfaction.

**Figure 3:**
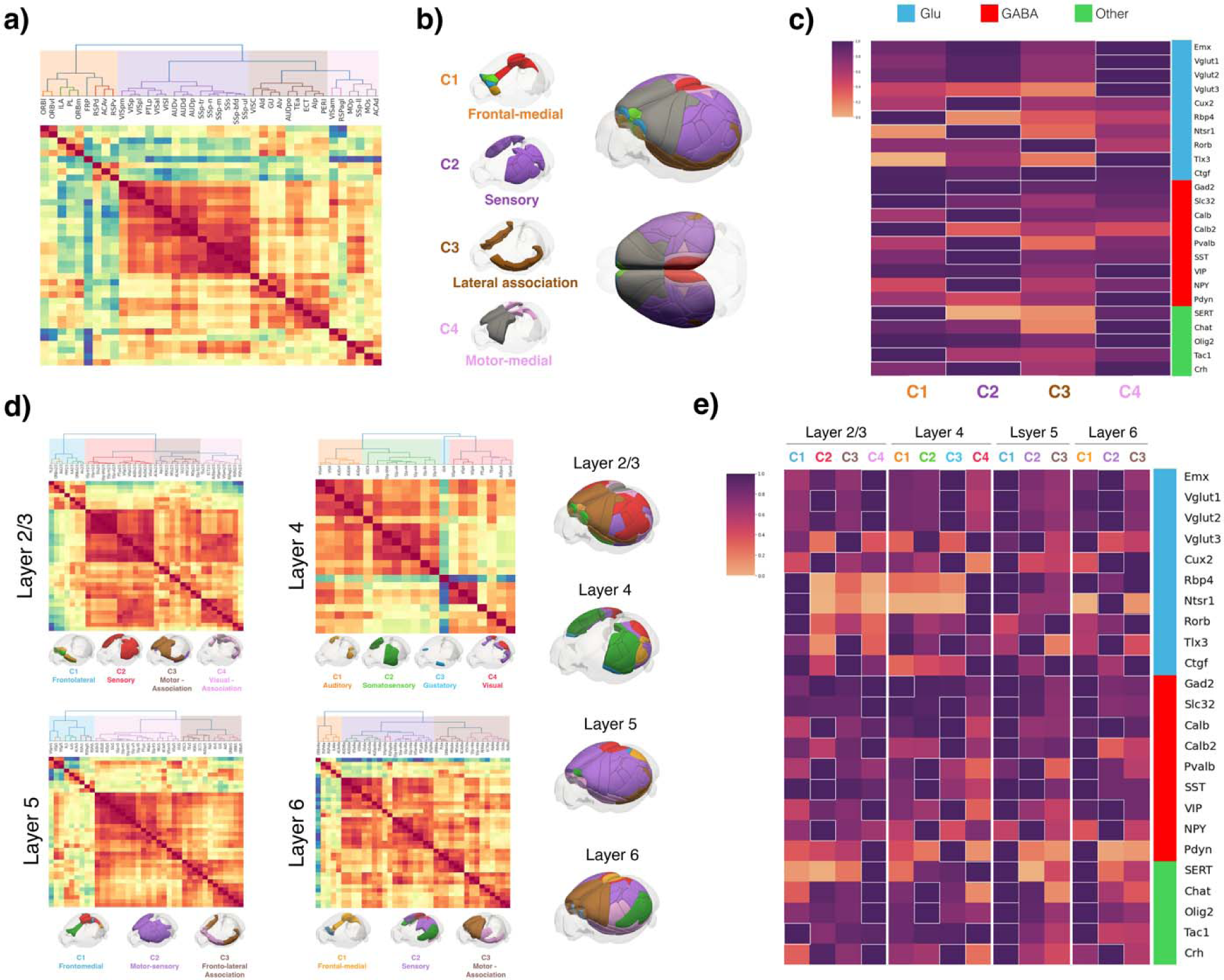
Cell type-based cortical organization. **a)** Cell type-based clustermap of the entire cortex revealing its organization into four major clusters, further subdivided into eight subclusters. **b)** Whole cortex 3D renderization showing the spatial distribution of all cortical areas according to the major cluster organization. Each individual area is color-coded based on the eight subclusters identified in the dendrogram (A). **c)** Normalized cell type distribution of major cortical clusters. The X-axis shows the major cortical clusters defined in A); the Y-axis shows all cell types clustered by normalized expression. The white boundaries delineate cell types with a higher normalized expression in specific area clusters. Cluster 1 is characterized by a high relative density of the GABAergic markers Slc32 and Calb2, the glutamatergic marker Rbp4, the serotoninergic marker SERT, and the neuropeptide Tac1. Cluster 2 is characterized by a high relative density of the GABAergic markers Gad2, NPY, Pv, Sst, and Calb, the glutamatergic markers Cux2 and Ntsr1, and the corticotropin-releasing factor Crh. Cluster 3 is characterized by a high relative density of the glutamatergic markers Rorb and Ctgf. Cluster 4 is characterized by a high relative density of the GABAergic markers VIP and Pdyn, the glutamatergic markers Emx, VGlut1, VGlut2, VGlut3, and Tlx3, the cholinergic marker Chat, and the oligodendrocytes marker Olig2. **d)** Layer-based cell type-based clustermaps showing distinct cortical organizations for each of the four cortical layers and the spatial organizations. Layer 2/3 shows four major clusters further subdivided into seven clusters. The lower panel shows the spatial distribution of each major cluster color-coded by the seven subclusters identified in the dendrogram. Layer 4 shows four major clusters further subdivided into five clusters. The lower panel shows the spatial distribution of each major cluster color-coded by the five subclusters identified in the dendrogram. Layer 5 shows three major clusters further subdivided into seven clusters. The lower panel shows the spatial distribution of each major cluster color-coded by the seven subclusters identified in the dendrogram. Layer 6 shows 3 major clusters further subdivided into seven clusters. The lower panel shows the spatial distribution of each major cluster color-coded by the seven subclusters identified in the dendrogram. The right panel shows the 3D renderization of each layer cluster organization with all clusters together. **e)** Normalized cell type distribution per layer of all layer-based clusters from D). The X-axis shows the major cortical clusters defined in D) for each layer; the Y-axis shows all cell types clustered by normalized expression. Cell-type clustermap divides cells cortical distribution into four main clusters. Layer 2/3 Cluster 1 is characterized by a high relative density of the glutamatergic markers Ntsr1, Rorb, Rbp4, and the tachykinin-releasing marker Tac1; Cluster 2 by a high relative density of the glutamatergic markers VGlut1 and Cux2, and the GABAergic markers NPY, Pv, Calb, and Sst; Cluster 3 by a high relative density of the glutamatergic markers VGlut3, Tlx3 and Ctgf; and Cluster 4 by a high relative density of the glutamatergic markers Emx and VGlut2, the GABAergic markers Gad2, Slc32, Calb2, Pdyn and VIP, the serotoninergic marker SERT, the cholinergic marker ChAT, the corticotropin-releasing hormone marker Crh, and the oligodendrocyte marker Olig2. Layer 4 Cluster 1 is characterized by a high relative density of the GABAergic marker Gad2; Cluster 2 by the GABAergic markers NPY, Sst and Pv, the cholinergic marker ChAT, and the oligodendrocyte marker Olig2; Cluster 3 by the glutamatergic markers Emx, VGlut1, VGlut2, Cux2 and Tlx3, the GABAergic markers Slc32, Pdyn, Calb, Calb2 and VIP, and the corticotropin-releasing hormone marker Crh; and Cluster 4 by the glutamatergic markers Ntsr1, Rorb, Rbp4, Ctgf and VGlut3, the serotoninergic marker SERT, and the tachykinin-releasing marker Tac1. Layer 5 Cluster 1 is characterized by a high relative density of the glutamatergic markers Emx, VGlut1, VGlut2, Cux2, Ntsr1, Rbp4 and Ctgf, the serotoninergic marker SERT, the GABAergic markers Gad2, Slc32, Pdyn, Calb, Calb2 and Sst, the tachykinin-releasing marker Tac1, the corticotropin-releasing hormone marker Crh, the cholinergic marker ChAT, and the oligodendrocyte marker Olig2; Cluster 2 by the glutamatergic markers VGlut3 and Tlx3, and the GABAergic markers NPY, Sst, and Pv; and Cluster 3 by the glutamatergic markers Ctgf and Rorb. Layer 6 Cluster 1 is characterized by a high relative density of the glutamatergic markers Rbp4 and VGlut3, the GABAergic markers Gad2, Slc32, Pdyn, Sst, Pv and Calb2, the serotoninergic marker SERT, the cholinergic marker ChAT, the tachykinin-releasing marker Tac1, and the oligodendrocyte marker Olig2; Cluster 2 by the glutamatergic markers Emx, VGlut1, VGlut2, Cux2, Ntsr1 and Tlx3, the GABAergic markers Calb, NPY and VIP, and the corticotropin-releasing hormone marker Crh; and Cluster 3 by the glutamatergic markers Ctgf and Rorb.

Upon closer examination, the regionally diverse Cluster 3 also reveals shared cell type compositions characterizing distinct subclusters with common functional specializations. Cluster C3.1 consists of the periaqueductal gray (PAG) and its interconnected regions, which integrate sensory, motor, and autonomic functions. This subcluster includes the midbrain reticular nucleus (MRN), cuneiform nucleus (CUN), midbrain reticular nucleus retrorubral area (RR), sensory-related medulla (MY-sen), and lateral habenula (LH). Cluster C3.2 comprises regions associated with the limbic system and basal ganglia circuits, including the striatum-like amygdalar nuclei, lateral septal complex, preparasubthalamic complex (PST), and subgeniculate nucleus (SubG). These areas are involved in motivation, emotion, and movement. Cluster C3.3 includes regions specialized in the processing and integration of auditory, olfactory, and upper limb somatosensory information. This subcluster contains the bed nucleus of the anterior commissure (BAC), inferior colliculus (IC), behavioral state-related medulla (MY-sat), midbrain trigeminal nucleus (MEV), and olfactory bulb areas. Cluster C3.4 is composed of regions involved in motivation, reward processing, social behaviors, motor control, and stress regulation, forming a dopamine-modulated network. This subcluster includes the ventromedial and dorsomedial hypothalamus, caudoputamen (CP), nucleus accumbens (ACB), and olfactory tubercle (OT). Cluster C3.5 includes all cerebellar regions, along with key motor and dopaminergic centers such as the reticular thalamus (RT), substantia nigra pars reticulata (SNr), substantia nigra pars compacta (SNc), ventral tegmental area (VTA), and red nucleus (RN). The SNr, SNc, and VTA are primary sources of dopaminergic neurons, which play crucial roles in movement, behavior, and reward processing. Along with cerebellar areas, these regions highlight the similar cellular organization of non-cortical motor centers. Cluster C3.6 contains a more heterogeneous set of regions, including the pons, motor-related medulla (MY-mot), dorsal pallidum (PALd), medial septal complex (MSC), substantia innominata (SI), medial habenula (MH), parabigeminal nucleus (PBG), oculomotor nucleus (III), pedunculopontine nucleus (PPN), and fundus striatum (FS). These regions are primarily associated with motor control and sensory-motor integration (pons, MY-mot, PALd, PPN, PBG, and III), with additional roles in attention and cognitive processing (PPN, MSC, SI, and MH). Together, these subclusters within Cluster 3 integrate sensory, motor, autonomic, emotional, and cognitive processes. Each subcluster highlights a distinct functional role, including sensory-motor integration, motivation, reward processing, and motor control.

### Cell type distribution can further define cortical region and layer organization

To further investigate cell type-based signatures across cortical regions and layers, we analyzed 24 selected cell types with cortical expression using similar hierarchical clustering of their densities as done for the whole brain above. We also visualized their distribution using flat-map-based cell density representations and representative example images (Figure 3, Supplementary Figure 1). First, we clustered cell type distributions across all cortical layers to generate an overview of their distinctions across different cortical areas. This analysis revealed four major cortical clusters and eight subclusters (Figure 3a-b, Methods): Cluster C1 includes frontomedial association cortical areas, further divided into four subclusters: C1.1: orbitolateral cortex, C1.2: prelimbic and infralimbic and orbitomedial cortex, C1.3: frontal pole and C1.4: anterior cingulate and retrosplenial cortical regions. Cluster C2 consists of somatosensory, visual, and auditory areas, and the posterior parietal association cortex, which serves as a connection hub between sensory areas and other cortical association region ^35–37^. Cluster C3 is primarily composed of lateral association areas. Cluster C4 consists mainly of somatomotor and medial association-related areas and can be further divided into two subclusters: C4.1: visual-association subcluster and C4.2: somatomotor medial subcluster.

Next, layer-specific analyses revealed overall shared cortical organization with the global cortical analysis with some distinctions. The fontal and medial C1 cluster organization is similarly displayed in layers 2/3, 5 and 6, with additional visual areas (VISp, VISpm and VISpl) included in the layer 5 C1^L5^ cluster (Figure 3c). The sensory and PTLp cluster C2 is also largely conserved across the layers 2/3, 5 and 6, with the additional inclusion of the motor areas in the layer 5 C2^L5^ cluster. The C3 lateral association cluster is similar in the layer 5 C3^L5^ cluster, while The C2 and C3 motor and lateral and medial association clusters are combined in the layer 2/3 C3^L2/3^ cluster and layer 6 C3^L6^. Finally, the analysis of layer 4 cell type distribution identified four major clusters across the granular cortex (Figure 3d). The cluster C1^L4^ is composed of mostly auditory areas, the cluster C2^L4^ corresponds to somatosensory areas and the visceral cortex (VISC), the cluster C3^L4^ comprises solely the gustatory cortex (GU), a multisensory integration area^38^, and the cluster C4^L4^ corresponds to mostly visual cortical areas. This organization reflects the functional specialization present within layer 4, wherein various sensory modalities exhibit distinct organization.

### Density-based hierarchical analysis of whole-brain cell type distribution

The above analyses used the anatomical atlas to investigate the cell type distribution, identifying shared cell type compositions for anatomical areas with common functions. This suggests that brain spatial organization may be examined in an unbiased manner by using cell type distribution patterns and local cell type densities as guiding factors.

To test this hypothesis, we analyzed the spatial distribution of each cell type across the whole brain using the DBSCAN (Density-Based Spatial Clustering of Applications with Noise) algorithm^39,40^ (Figure 4a). First, we chose ε = 150 um to define the radius around each cell within which the local cell densities were calculated. Second, for each detected cell nucleus, we counted the number of neighboring cells within this radius and generated a histogram showing the frequency distribution of cell densities for each cell type. And third, we divided each cell type’s density distribution into nine percentiles, ranging from the 10th to the 90th percentile, where the nth percentile represents the density threshold below which n% of the cell densities fall. For example, the 10th percentile density for Cux2-expressing cells corresponds to 49,797 cells/mm³, identifying broad spatial areas with Cux2 densities above this low threshold (Figure 4b). In contrast, the 90th percentile density for Cux2-expressing cells corresponds to 529,455 cells/mm³, identifying discrete spatial areas with Cux2 densities above this high threshold (Figure 4b).

**Figure 4:**
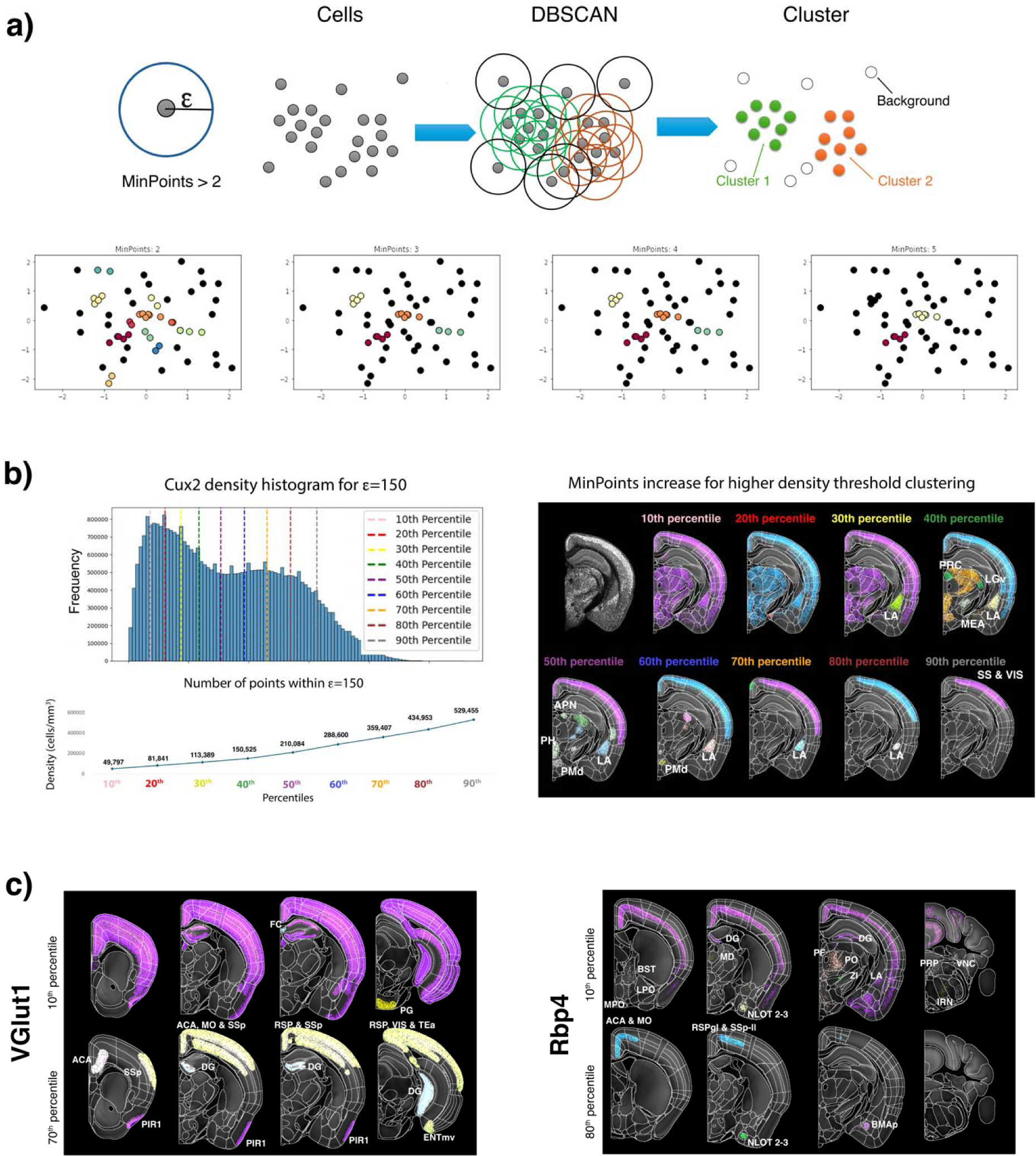
Density-based whole brain spatial organization. **a)** Scheme showing density-based DBSCAN spatial clustering of brain cell distribution. ε defines the maximum distance (radious) between two points for them to be considered neighbors (in our analysis ε=150 μm); MinPoints defines the minimum number of points/neighbors within ε requiered to be considered a dense region, i.e. a cluster. Lower panel shows how the change in the number of MinPoints affects the number of clusters in a small synthetic dataset. **b)** The definition of MinPoints is based on the histogram density frequency (left panel), thus taking into account the varying density of each individual cell type based on its intrinsic cell density distributions. MinPoints values are selected using 9 percentiles from the histogram density distribution (from 10^th^ to 90^th^ percentile), in this example for Cux2. Afterwards, it is analyzed nine different DBSCAN clusterings, one for each percentile threshold, and the resulting spatial density clusters are individually visualized in the whole brain (right panel), with each color representing the cells belonging to an individual cell density cluster. **c)** Coronal planes showing the spatial-density distribution at 10^th^ percentile and 70^th^ percentile for VGlut1 and 80^th^ percentile for Rbp4. Coronal planes are overlaid onto the CCF reference brain and the CCFv3 atlas to show the anatomical distributions of the density analysis.

The examination of the Cux2 density distribution across the nine percentiles indeed revealed partitioning of distinct anatomical areas at different Cux2 cell densities (Figure 4b). While the 10^th^ percentile highlights broad cortical areas, starting with the 30^th^ percentile, distinct cortical and subcortical subregions become delineated. For example, the lateral amygdala (LA) becomes spatially defined as an individual spatial cluster by the 30^th^ percentile density of 113,389 cells/mm3, while the LA ventral part is further delineated by the 80^th^ percentile density of 434,952 cells/mm3. This unbiased LA delineation aligns well with expert-defined anatomical boundaries based on Nissl staining, immunohistochemistry ^41^ and tract-tracing studies ^42^. Other areas, such the precommisural nucleus (PRC), ventral part of the lateral geniculate (LGv), and medial amygdala (MEA) are highlighted as individual spatial clusters by the 40^th^ percentile Cux2 density of 150,525 cells/mm3, the anterior pretecta nucleus (APN), posterior hypothalamic nucleus (PH) and the dorsal premammillary nucleus (PMd) are highlighted by the 50^th^ percentile density of 210,084 cells/mm3, and superficial layers of the somatosensory (SS) and visual (VIS) cortices are highlighted by the 90^th^ percentile density of 529,455 cells. Thus the analysis of a single Cux2+ cell type reveals that distinct cell densities are associated with and outline the spatial borders of distinct anatomical areas, effectively partitioning brain regions and their subregions and into a cell density-based hierarchical organization.

Next, we applied this approach to analyze the cortical distribution of glutamatergic pyramidal neurons identified by VGlut1 and layer 5-specific Rbp4 expression (Figure 4c). The cell density segmentation of the VGlut1 distribution revealed a broad cortical pattern at the 10^th^ percentile density (202,445 cells/mm3), while the 70th percentile density (395,836 cells/mm³) selectively highlighted the frontomedial, motor, and lateral association areas and sensory areas. Similarly, the Rbp4-expressing neurons followed the expected layer 5 distribution at the 10th percentile density (25,606 cells/mm³), but by the 50th percentile density (169,836 cells/mm³), the segmentation distinctly emphasized the frontomedial and motor cortical areas. Consistent with the findings from Cux2 density analysis, the spatial distribution of VGlut1 and Rbp4 reveals a density-based gradient that highlights cortical regions with shared functions, reinforcing the concept of cell density-based hierarchical organization of cortical areas. A full cell density-based segmentation of cortical and subcortical areas, based on the Cux2, VGlut1 and Rbp4 cell types, is shown in Supplementary Movies 5-7.

The caudoputamen, part of the striatum, plays a crucial role in motor control, habit formation, and reinforcement learning, integrating sensorimotor, cognitive, and reward-related inputs which can be used to spatially subdivide the CP into different functional domains^4,43^. The main cell type in the CP are the medium spiny neurons, which are further subdivided into the D1 and D2 dopamine receptor-expressing subpopulations. Here, we analyzed the caudoputamen spatial cell type distribution using three gene markers—Gad2 marking all medium spiny neurons, Calb marking the D2-positive and Pdyn marking the D1-positive medium spiny neurons. This analysis revealed that their cell densities uniquely highlight distinct CP subdivisions (Figure 5). Within the intermediate CP (Figure 5a), the ventrolateral region of the medial part of the CP is broadly delineated by Calb 80^th^ percentile density (402,202 cells/ mm3), corresponding to the ventral and medial striatal domains, which receive inputs from the sensory and lateral association regions. These same regions are delineated by Gad2 70th percentile density (272,473 cells/ mm^3^), while the highest density of Gad2, the 90th percentile (382,467 cells/mm^3^), highlights a distinct subregion receiving inputs from the inner-mouth sensorimotor cortex, corresponding to the ventrolateral striatal domain, a subdivision associated with the inner mouth sensory-motor network. Similarly, Pdyn cell density at 60th and 80th percentiles (246,937 cells/ mm^3^ and 353,111 cells/ mm^3^, respectively) delineates the sensory and lateral association regions of the CP, although circumscribed to the ventrolateral, ventromedial and dorsomedial domains.

**Figure 5:**
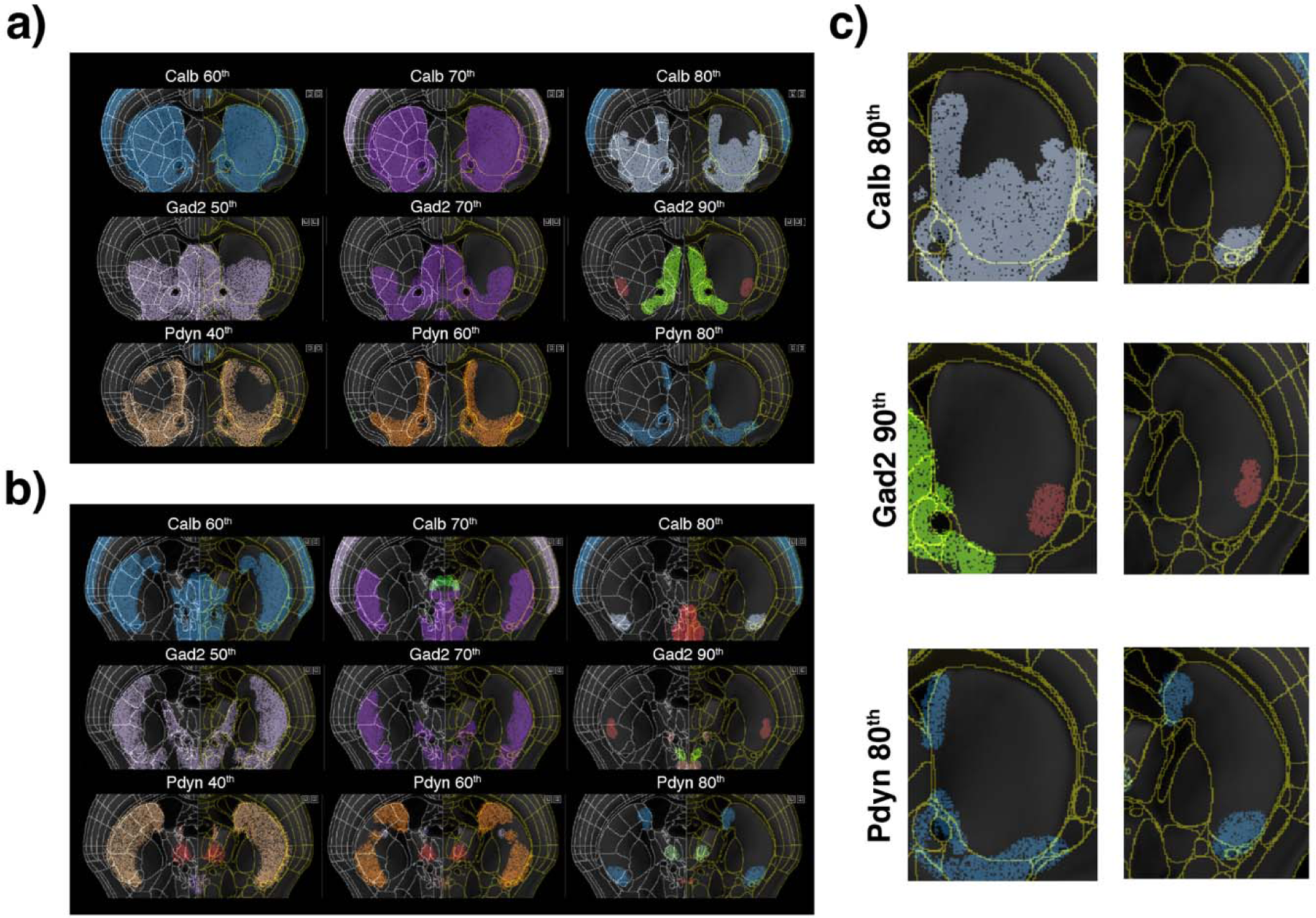
Density-based spatial organization of the CP. **a)** Coronal planes showing th density-based cluster analysis of Calb, Gad2 and Pdyn in the intermediate CP. At the 60^th^ percentile density Calb delineates the entire CP, however at 80^th^ percentile it delineates subregions corresponding to the striatal domains Cpi.vl.vt (m/o), CPi.vl.v (m/i), CPi.vl.imv (ul), CPi.dl.imd (II), CPi.vm.v, CPi.vm.vm, CPi.vm.cvm, CPi.vl.cvl, CPi.dm.im, and CPi.dm.dm. On the other hand, Gad2 at the 90th percentile highlights the striatal domain CPi.vl.v (m/i), which receives inputs from the inner-mouth sensorimotor cortex. Similarly, Pdyn delineates the sensory and lateral association regions of the CP, but at 80^th^ percentile it is restricted to the striatal subdomains Cpi.vl.vt (m/o), CPi.vl.v (m/i), CPi.vm.v, CPi.vm.vm, CPi.dm.im, and CPi.dm.dm. **b)** Coronal planes showing the density-based cluster analysis in the caudal CP. All three cell- types delineate the entire CP a the lower densities analyzed. However, Gad2 and Calb at 70^th^ percentile, Pdyn at 60^th^ percentile delineate the striatal domains CPc.v.m, CPc.v.dl, CPc.i.vl and CPc.i.dl. Pdyn also delineates at 60^th^ percentile the striatal domains CPc.d.dm and CPc.d.dl. Calb and Pdyn, both at 80^th^ percentile, delineate the striatal ventral domain CPc.v.m, and Pdyn also delineates the caudal domain CPc.d.dm. Gad2 at 90^th^ percentile delineates de ventral striatal domain CPc.v.dl. **c)** Details of Calb, Gad2 and Pdyn density-cluster analysis showing CP delineation at higher densities in both the intermediate (left column) and caudal (right column) CP.

In the more caudal CP (Figure 5b), the Calb at the 70^th^ percentile, Gad2 at the 70^th^ percentile, and Pdyn at the 60^th^ percentile densities highlight the ventral and intermediate striatal domains receiving inputs from lateral association areas. Pdyn 60^th^ percentile density further delineates the dorsal striatal domains receiving cortical medial connections. At the higher 80^th^ percentile densities, both Calb and Pdyn delineate the striatal centroventral medial domain, receiving inputs from the NAcc shell, as well as the caudal dorsomedial domain receiving inputs from the dorsomedial SNr, thus integrating motor and association functions. Finally, the highest Gad2 cell density, the 90^th^ percentile, delineates the caudal ventral striatal domain, integrating inputs from anterolateral SNr. Taken together, similar to the cortical data, the distribution of major CP cell types reveals a density-based hierarchical spatial organization linked to different CP functional domains, as defined by their input-output relationships.

Next, we extended the cell density analysis to major subcortical regions. The neuropeptide NPY, VGlut2, Rbp4 and PV cell densities highlight the thalamus and its different subdivisions. For example, the NPY 10^th^ percentile (54,961 cells/mm^3^) selectively highlights several thalamic nuclei, including the anteroventral nucleus of the thalamus (AV), the ventral anterior-lateral plex of the thalamus (VAL), the ventral medial nucleus of the thalamus (VM), the nucleus reuniens (RE), the reticular thalamus (RT), the ventral posteromedial nucleus of the thalamus (VPM), the medial habenula (MH), the paraventricular nucleus of the thalamus (PVT), the parafascicular (PF) and the ventral part of the lateral geniculate (LGv), while the RT is highlighted by the 40^th^ percentile (159,508 cells/mm^3^). These regions are involved in both sensoriomotor (VPM, VM, VAL, LGv) processing and limbic-memory functions (AV, RE, PVT, MH), while RT plays a key role in modulating thalamocortical activity.

Most regions of the thalamus, with the exception of the reticular thalamus, are highlighted by the VGlut2 cell density distribution, while the high 80^th^ percentile VGlut2 density (481,567 cells/mm^3^) further delineates the parafascicular nucleus (PF), nucleus reuniens (RE), central medial nucleus of the thalamus (CM), interanteromedial nucleus of the thalamus (IAM), and rhomboid nucleus (RH) (Figure 6). These regions are all part of the midline and intralaminar thalamic nuclei, involved in memory, decision-making, and goal-directed behavior, with strong projections to the prefrontal cortex, basal ganglia, and hippocampus. The mediodorsal (MD), the suprafascicular, magnocellular part (SPFm), parafascicular (PF) and the posterior complex (PO) nuclei of the thalamus are highlighted by Rbp4 expression at 10^th^ percentile density (25,606 cells/mm^3^), with the PF and the medial PO being further highlighted by the 30^th^ percentile density (104,900 cells/mm^3^). These thalamic nuclei are part of the limbic midline thalamus and have been previously shown to have strong cortical and basal ganglia connections^44^. Pv at 10^th^ percentile density (19,947 cells/mm^3^) delineates the RT and the LGv, with subregions of RT further delineated at the 40^th^ percentile density (114,874 cells/mm^3^). Both regions are reciprocally connected, creating a feedback loop that modulates sensory processing and attention^45^, highlighting the importance of Pv in general sensory processing.

**Figure 6:**
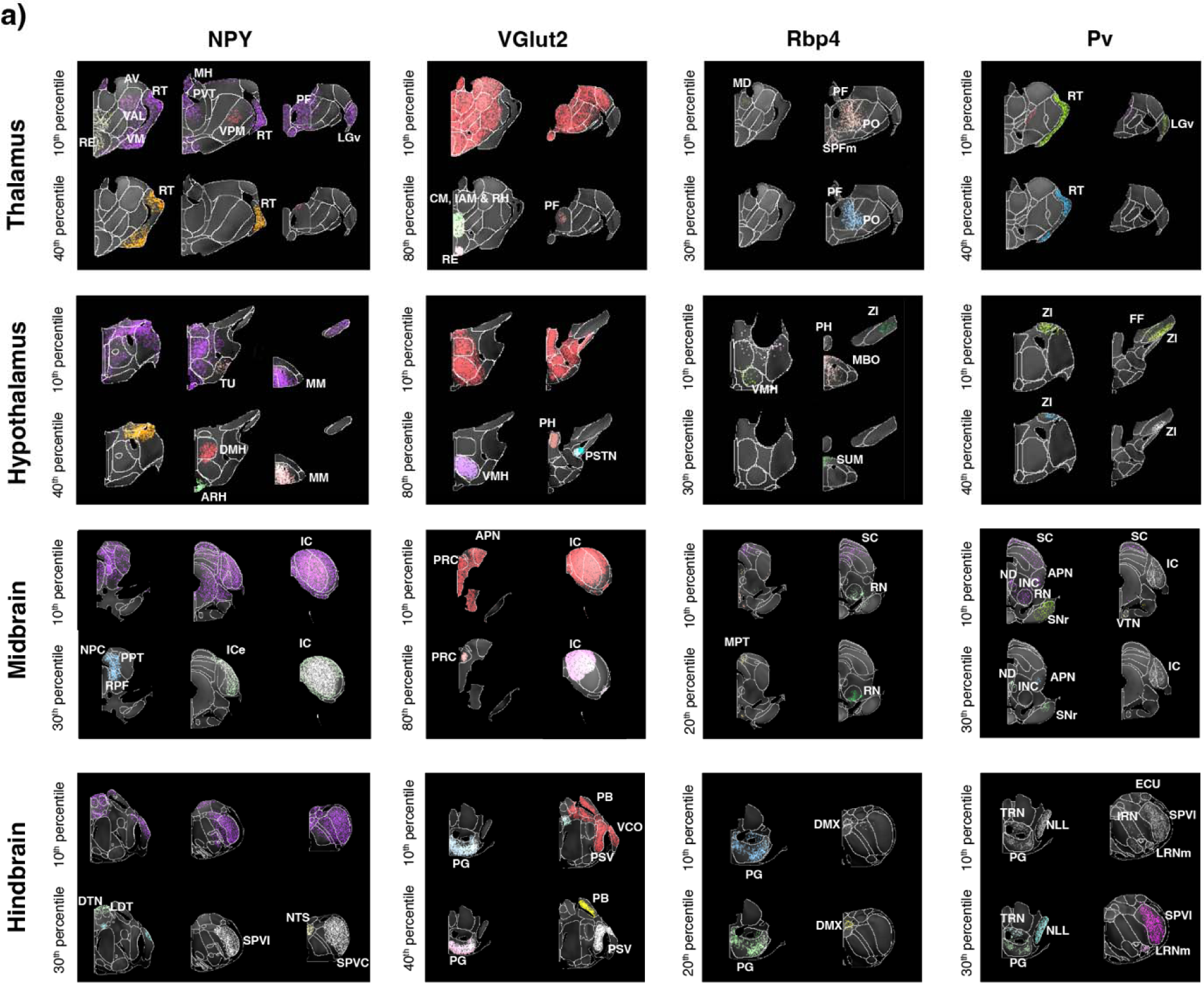
Density-based spatial organization in the major subcortical regions. **a)** Density-based cluster analysis of NPY, VGlut2, Rbp4 and PV in the thalamus, hypothalamus, midbrain and hindbrain. Cell density patterns are visualized at specific percentile thresholds to reveal the spatial clustering and regional specificity of each cell population. NPY is displayed at 10th, 40th percentiles (Thalamus and Hypothalamus) and 10th, 30th percentiles (Midbrain and Hindbrain); VGlut2 at 10th, 50th percentiles (Thalamus, Hypothalamus, and Midbrain) and 10th, 40th percentiles (Hindbrain); Rbp4 at 10th, 30th percentiles (Thalamus and Hypothalamus) and 10th, 20th percentiles (Midbrain and Hindbrain); and Pv at 10th, 40th percentiles (Thalamus and Hypothalamus) and 10th, 30th percentiles (Midbrain and Hindbrain). All datasets cluster analyses are overlay onto the CCF reference brain and the CCFv3 anatomical atlas to outline anatomical boundaries of nuclei and subnuclei. In the Thalamus, NPY delineates the reticular nucleus (RT) and ventral posteromedial nucleus (VPM), while Pv neurons delineates the RT. In Hypothalamus, NPY neurons highlights the tuberal nucleus (TU) and ventromedial hypothalamic nucleus (VMH), with both NPY and VGlut2 populations present in the arcuate hypothalamic nucleus (ARH). The Midbrain shows prominent Rbp4 expression in the red nucleus (RN) and both Rbp4 and Pv expression in the inferior colliculus (IC). In the Hindbrain, VGlut2 neurons are enriched in the parabrachial nucleus (PB), while the pontine gray (PG) contains populations of multiple cell types, and NPY neurons are detected in the nucleus of the solitary tract (NTS). *Thalamus*: LGv, Lateral geniculate nucleus, ventral part; MD, Mediodorsal nucleus; PF, Parafascicular nucleus; RE, Nucleus of reuniens; RT, Reticular nucleus; SPFm, Subparafascicular nucleus, magnocellular part; VPM, Ventral posteromedial nucleus. *Hypothalamus*: ARH, Arcuate hypothalamic nucleus; DMH, Dorsomedial nucleus of the hypothalamus; MBO, Mammillary body; MM, Medial mammillary nucleus; PH, Posterior hypothalamic nucleus; PSTN, Parasubthalamic nucleus; SUMm, Supramammillary nucleus, medial part; TU, Tuberal nucleus; VMH, Ventromedial hypothalamic nucleus; ZI, Zona incerta. *Midbrain*: APN, Anterior pretectal nucleus; IC, Inferior colliculus; MPT, Medial pretectal area; NPC, Nucleus of posterior commissure; PRC, Precommissural nucleus; RN, Red nucleus; SCm, Superior colliculus, motor related; SNr, Substantia nigra, reticular part; VTN, Ventral tegmental nucleus. *Hindbrain*: DMX, Dorsal motor nucleus of the vagus; DTN, Dorsal tegmental nucleus; ECU, External cuneate nucleus; LRNm, Lateral reticular nucleus, magnocellular part; NLL, Nucleus of the lateral lemniscus; NTS, Nucleus of the solitary tract; PB, Parabrachial nucleus; PG, Pontine gray; PSV, Principal sensory nucleus of the trigeminal; SPVi, Spinal vestibular nucleus, inferior part; SPVl, Spinal vestibular nucleus, lateral part; TRN, Tegmental reticular nucleus; VCO, Ventral cochlear nucleus

Multiple hypothalamic regions are also highlighted by specific cell type densities. NPY 10^th^ percentile density highlights multiple broad medial hypothalamic regions as well as the tuberal nucleus (TU) as a single spatial cluster, while the NPY 40^th^ percentile density highlights the dorsomedial nucleus of the hypothalamus (DMH), the arcuate hypothalamic nucleus (ARH), and the medial mammillary nucleus (MM) which together contribute to regulating homeostasis (TU, ARH and DMH) and stress (DMH and MM). VGlut2 highlights a large portion of the hypothalamus at the 10^th^ percentile density, while the 80^th^ percentile density further delineates the ventromedial hypothalamus (VMH), posterior hypothalamic nucleus (PH) and the parasubthalamic nucleus (PSTN) connected with the limbic system and brainstem. The Rbp4 at 10^th^ percentile density highlights the VMH, the PH, the zona incerta (ZI), the supramammillary nucleus (SUM), and the mammillary body (MBO), while at 30^th^ percentile density it highlights selectively the medial part of the SUM. These nuclei integrate limbic, hypothalamic, and brainstem functions, with the SUM nucleus playing essential roles hippocampal functions^46^. Pv at 10^th^ percentile density delineates the interconnected ZI and the Fields of Forel (FF), with a subregion of ZI further delineated at the PV 40^th^ percentile density, acting as the primary connection between the striatopallidal and the thalamocortical network^47^. Moreover, the higher Pv cell density in the ZI defines the ZIc (ZI caudal part), a subregion of ZI previously shown to be connected primarily with the midbrain and the pons^48^.

The structures of the midbrain and hindbrain can also be uniquely highlighted by specific cell types (Figure 6). Broad definition of the midbrain and hindbrain areas is delineated by NPY at the 10^th^ percentile, while the nucleus of the posterior commissure (NPC), the posterior pretectal nucleus (PPT), the retroparafascicular nucleus (RPF), the dorsal tegmental nucleus (DTN), the laterodorsal tegmental nucleus (LDT), the spinal nucleus of the trigeminal, interpolar part (SPVI), the nucleus of the solitary tract (NTS), and the spinal nucleus of the trigeminal, caudal part (SPVC) are further delineated by the NPY 30^th^ percentile (118,411 cells/mm^3^). VGlut2 delineates areas in both structures across multiple percentile densities. The inferior colliculus (IC), the precommisural nucleus (PRC), the anterior pretectal nucleus (APN), the pontine grey (PG), the parabrachial nucleus (PB), the ventral cochlear nucleus (VCO), and the principal sensory nucleus of the trigeminal (PSV) are all highlighted by the VGlut2 10^th^ percentile, while the hindbrain PG, the PSV and the PB are highlighted by the 40^th^ percentile (327,010 cells/mm^3^) and the PRC and the IC are further highlighted by the 80^th^ percentile density. The IC, VCO, and PB are primarily involved in auditory processing, while the PSV, APN, and PB process somatosensory and pain signals. The PG and PRC contribute to motor coordination and sensory-motor integration. Altogether, these regions are primarily related to sensory function, indicating the influence of glutamatergic signaling in sensory processing. The medial pretectal area (MPT), the superior colliculus (SC), the red nucleus (RN), the PG, and the dorsal motor nucleus of the vagus nerve (DMX) are all highlighted by Rbp4 at the 10^th^ percentile density, while the 20^th^ percentile (60,125 cells/mm^3^) further highlights the MPT, RN, PG and DMX. These regions work together to regulate autonomic functions, coordinate motor control, and especially sensorimotor integration (MPT, RN, PG and DMX). Pv has a broad expression within the midbrain and the hindbrain, delineating at 10^th^ percentile density the SC, IC, nucleus of Darkschewitsch (ND), interstitial nucleus of Cajal (INC), red nucleus (RN), ventral tegmental nucleus (VTN), and substantia nigra pars reticula (SNr) in the midbrain and SPVI, PG, tegmental reticular nucleus (TRN), nucleus of the lateral lemniscus (NLL), and lateral reticular nucleus magnocellular part (LRNm). At 30^th^ percentile Pv further delineates the ND, INC, APN and SNr within the midbrain, and the TRN, PG, NLL, SPVI and LRNm within the hindbrain. Previous works have demonstrated that APN, RN, and SNr are involved in the regulation of motor movements. The SNr serves as the output portal of the basal ganglia, playing a crucial role in the modulation of motor activity. The higher density within the SNr further delineates the SNr.oro-brachial domain, which receives inputs from the striatal domain that are heavily innervated by the somatic sensorimotor cortex in controlling orofacial movements^43,49^. The APN, which receives inputs from the SC, is primarily responsible for controlling eye movements. Meanwhile, the RN generates direct projections to the spinal cord, contributing to the coordination of motor functions. Together, these structures form an intricate network that ensures precise and coordinated motor control.

The hippocampal formation comprises the dentate gyrus (DG), CA3, CA2, and CA1 subregions of the main hippocampal circuit, as well as the fasciola cinereum (FC), a distinct subregion projecting to the DG and proposed to play a critical role in visual contextual memory^50^. Strikingly, all these areas can be spatially defined by distinct densities of a few cell types (Figure 7a). For instance, the DG granule cell layer (DG-sg) is delineated by Calb2 at the 60^th^ percentile density (270,139 cells/mm3) and VGlut1 at the 10th percentile (202,445 cells/mm3). The CA3 (CA3i) is defined a gap in cell clustering due to lower cell densities than the VGlut1 10th percentile (density 202,445 cells/mm3) or Calb2 20^th^ percentile (124,990 cells/mm3). Additional subregion of the CA3, the CA3dd, is defined by the serotoninergic marker SERT 30^th^ percentile (187,308 cells/mm3). The CA1 is delineated by VGlut2 at the 30th percentile density (288,176 cells/mm3). As illustrated in the right diagram (Figure 7a), the overlay of delineations from the preceding markers facilitates the definition of the CA2 region, situated between the clusters that delineate the CA1 and CA3i regions. Finally, the FC is delineated by the calcium-binding protein Calb2 at the 40^th^ percentile density (169,694 cells/mm3), the glutamatergic markers VGlut1 at the 10th percentile (202,445 cells/mm3) and VGlut2 20^th^ percentile (243,047 cells/mm3) densities, and the serotoninergic marker SERT at the 10^th^ percentile (52,274 cells/mm3).

**Figure 7:**
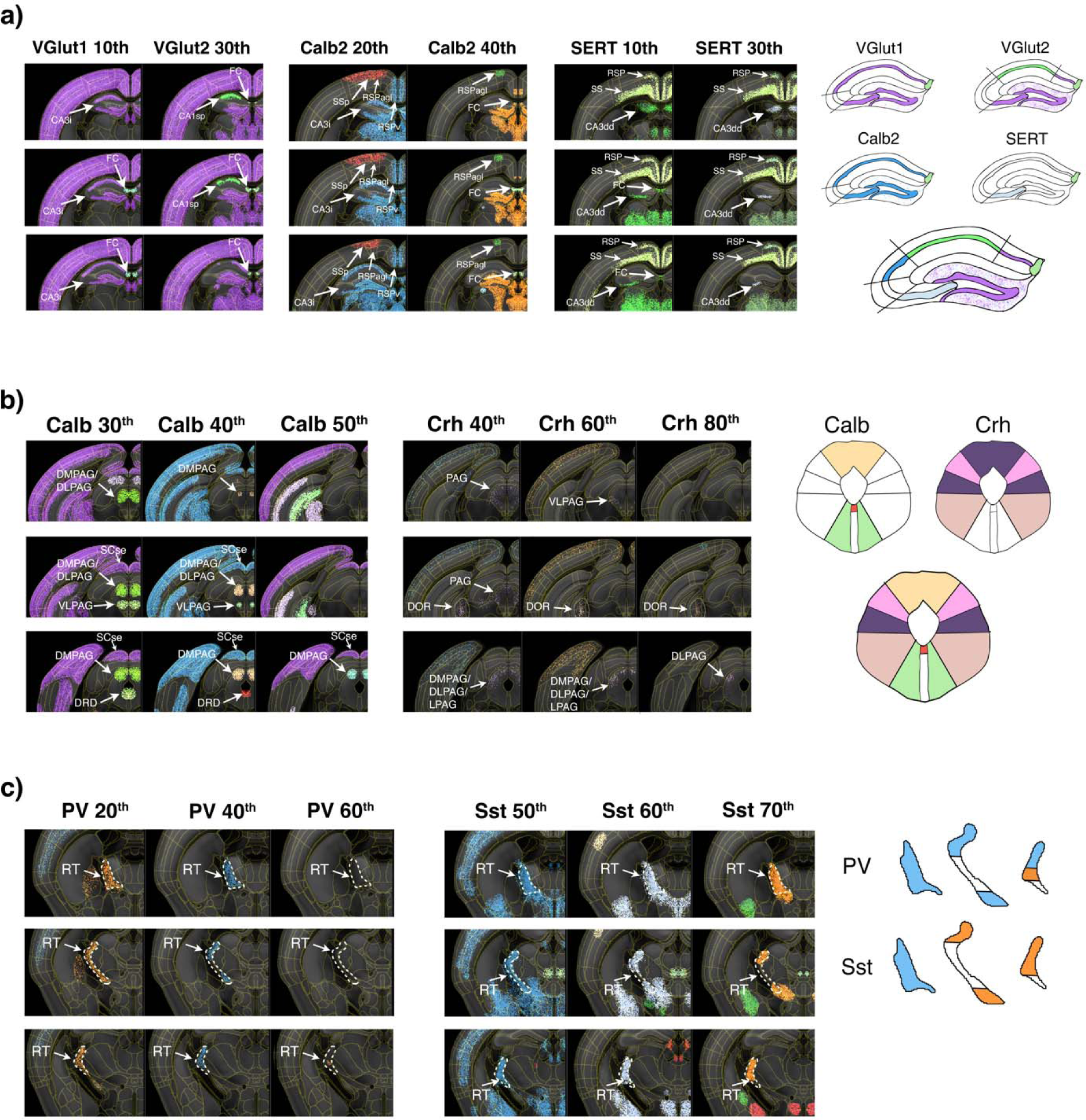
Density-based spatial organization of combined cell types. **a)** Coronal view of density-based cluster analysis for VGlut1 and VGlut2 (10^th^ and 30^th^ percentile thresholds), Calb2 (20^th^ and 40t percentile thresholds) and SERT (20^th^ and 30^th^ percentile thresholds) in the hippocampal formation. The right diagram shows the composition of the hippocampal formation delineations from both VGlut1, VGlut2, Calb2 and SERT. **b)** Density-based analysis of Calb (30^th^, 40^th^, and 50^th^ percentile thresholds) and Crh (40^th^, 60^th^, and 80^th^ percentiles) in the PAG. Delineated regions include DMPAG, DLPAG, LPAG, and VLPAG. The panel also shows clusters in raphe nuclei regions including DRD and DRV. The right diagram shows the composition of the PAG subdivisions from both Calb and Crh. **d)** Density-based analysis of PV (20^th^, 40^th^, and 60^th^ percentiles) and Sst (50^th^, 60^th^, and 70^th^ percentiles) clusters in the RT. The different thresholds show a spatial distribution of both Pv and Sst within the RT. The right diagram shows the composition of the RT subdivisions from both Pv and Sst. *Hippocampal Formation*: CA1, Cornu Ammonis 1; CA2, Cornu Ammonis 2; CA3, Cornu Ammonis 3; CA3dd, CA3 dorsal, dorsal part; CA3i, CA3 intermediate; CA3sp, CA3 pyramidal layer; DG, Dentate Gyrus, FC, Fascicula Cinerea. Periaqueductal Gray: DMPAG, Dorsomedial Periaqueductal Gray; DLPAG, Dorsolateral Periaqueductal Gray; LPAG, Lateral Periaqueductal Gray; VLPAG, Ventrolateral Periaqueductal Gray. Raphe Nuclei: DRD, Dorsal Raphe Nucleus, Dorsal Part. Thalamus: RT, Reticular Thalamic Nucleus.

The periaqueductal grey (PAG) is a structure surrounding the cerebral aqueduct, with known roles in autonomic function, motivated behavior, and responses to threatening stimuli^51,52^. PAG is typically subdivided into four major subdivisions based on distinct input/output connectivity underlying its distinct functions^53,54^. Applying the cell density-based spatial segmentation, we can identify all major subdivisions of the PAG by distinct cell densities of a few cell types (Figure 7b). The entire rostral PAG is delineated as a single cluster by Crh at the 40th percentile density (17,401 cells/mm3). The dorsomedial PAG (DMPAG) is delineated by Calb at the 30th and 40th percentiles (184,761 cells/mm3 and 229,112 cells/mm3, respectively), with the 50th percentile (272,332 cells/mm3) delineating only the more caudal part, and by Crh at the 40th and 60th (28,294 cells/mm3) percentiles, also delineating the more caudal part. The dorsolateral (DLPAG) is delineated by Calb at the 30th percentile in the rostral and medial parts, at the 40th percentile in the medial part, and by Crh from the 40th to the 80th (45,200 cells/mm3) percentiles. The lateral PAG (LPAG) is delineated by Crh at the 40th and 60th percentiles. The ventrolateral (VLPAG) is delineated by Crh at the 60th percentile and by Calb at the 30th percentile and at the 50th percentile only in the more medial part. The dorsal raphe (DRD) is delineated by Calb at the 30th and 40th percentiles.

The reticular nucleus of the thalamus (RT) comprises inhibitory GABAergic neurons which modulate thalamic responses to cortical inputs^55,56^. Previous studies have defined prominent RT cell types by the expression of Pv and Sst markers^57^. Our density-based analysis shows a clear spatial organization of these cell types within the RT (Figure 7c). The whole RT structure is delineated by Pv 20^th^ percentile (35,368 cells/mm3) and Sst percentile 50^th^ percentile (96,059 cells/mm3). However, the Pv 40^th^ percentile (114,875 cells/mm3) highlights the dorsal and ventral RT, without the medial RT section. Similarly selective dorsal and ventral, but not medial, segmentation is seen at Sst 60^th^ and even more prominent at 70^th^ percentile (121,807 and 197,847 cells/mm3, respectively). This segmentation corresponds well with established RT connectivity patterns: the dorsal part connects with visual areas, the ventral part with visceral and gustatory regions, and the medial part with somatosensory areas, while the posterior part of the medial RT connects with auditory regions^57–59^. Thus, the density-based cell distribution of Pv and Sst highlights previously defined topographical functional organization of the RT.

## DISCUSSION

In the current study we have characterized the 3D spatial distribution of 30 different cell populations defined by cell type-specific Cre mouse lines across the entire mouse brain at single cell resolution using a largely automated whole-brain analysis pipeline, including automated machine learning-based single-cell segmentation, a registration platform using intrinsic anatomical features (IAFs), and an unbiased cell density-based spatial distribution analysis and 3D spatial area parcellation.

We used these methods to study brain organization using both males and females. While most of these Cre lines correctly label the cell type populations expressing the respective cell-type marker gene in the adult brain, we also included four Cre lines driven by gene markers with strong developmental expression that is not sustain in the adult brain. This includes nNOS-Cre-based broad cell labeling across all brain regions^18^; SERT-Cre-based dense cell labeling in the cortex and the thalamus, reflecting the role of SERT in the formation of cortico-thalamic connections^21,22^, VGlut2-Cre-based cortical cell labeling^24–26^, and VGlut3-Cre-based cell labeling in distinct cortical regions and the thalamus. Since all these mice are knock-in Cre mice with Cre expression directly linked to the endogenous genes, the adult brain labeling reflects the adult cell lineages marked by the transient developmental expression, rather than artificial Cre misexpression. We therefore included these mouse lines in the current analysis, with the caveat of developmental contribution to the adult cell counts and distributions.

Our data first revealed that, while the numbers of overall neurons (Snap25) and total glutamatergic neurons (VGlut1) are similar between males and females, there are significant sex-specific differences in the profiles of glutamatergic neurons. Males exhibit increased levels of Cux2+, Rbp4+, Rorb+, Tlx3+ and VGlut2 neurons while females show higher VGlut3+ neurons. These changes in developmental glutamatergic markers without alterations in the overall number of glutamatergic neurons suggest a sex-dimorphism on cortical development related to neuronal differentiation, layer specification, and the establishment of connectivity patterns. Additionally, we also identified brain-wide larger numbers of Olig2+ cells in males, extending on previous findings of regional sex differences in myelin markers^33,34^. This pattern may underlie observed sex differences in white matter microstructure and potentially contribute to differential susceptibility to neurological disorders associated with oligodendrocyte dysfunction, such as multiple sclerosis^60^.

Next, we showed that major brain anatomical areas can be clustered and defined by their distinct cell-type composition signatures. While differences in cell type distribution across brain regions are well known and have been most recently demonstrated by a whole mouse brain MERFISH study, which identified as many as 300 cell types based on an analysis of more than 5000 gene transcript ^61^, our study demonstrates that a relatively few major cell type-specific gene markers in the brain are sufficient to cluster functionally related brain areas of the cortex, hippocampus, thalamus and hypothalamus and other brain areas. This data thus emphasizes the importance of these major cell type populations in defining the functional architecture of the brain.

Finally, given the power of the selected cell type markers in characterizing functionally related brain areas, we asked whether the distribution of individual cell types may be sufficient to identify brain regions and subregions with shares subregions. We tested this hypothesis using unbiased spatial clustering of the cell type cell densities by DBSCAN, with progressive segmentation of the whole-brain data from the 10^th^ to 90^th^ cell density percentile of each cell type. This novel approach revealed a cell density-based hierarchical organization of the brain, where broad regions are progressively subdivided into functionally specialized subregions based on distinct cell type cell densities. For example, all cortical areas are identified as a shared spatial cluster defined by low densities of cortical pyramidal neuron markers Cux2, VGlut1 and Rbp4, while higher thresholds densities selective highlight the frontomedial, motor, lateral association and sensory areas comprising relatively higher neuronal densities identified by these gene markers.

Similarly, Pv+ neuronal population marks broad spatial areas in the midbrain at low densities, while higher Pv+ cell densities identify a discrete regions, such as the APN, RN, and SNr involved in the regulation of motor movements, with the highest density marking the SNr.oro-brachial domain^49^. The cell density-based hierarchical organization is also seen in the analysis of the CP, which reveals that distinct cell-type cell densities delineate increasingly refined CP subregions that align with functional domains defined by CP inputs. For example, the ventral and medial CP domains are selectively highlighted by high densities of Gad2 (both D1+ and D2+ medium spiny neurons), Calb (D2+ medium spiny neurons) and Pdyn (D1+ medium spiny neurons) cell densities, revealing cell density-based distinctions between ventral and medial CP domains integrating inputs from the sensory and lateral association regions, including the a distinct subregion receiving inputs from the inner-mouth sensorimotor cortex highlighted by the Gad2 90th percentile of 382,467 cells/mm3, and lateral CP domains integrating motor-related inputs. Strikingly, all subregions of the hippocampal formation can be identified by cell density-based spatial partitioning of only 4 gene markers, Calb2, SERT, VGlut1 and VGlut2, contrasting with earlier methods that required profiling over 200 genes to identify these subregions in tissue sections^62,63^.

The DBSCAN approach extends beyond traditional anatomical studies, which rely on expert-based analysis of Nissl staining and a limited set of cyto- and immunolabeling methods, by leveraging intrinsic cellular architecture to define regions and subregions. Exploiting distinct cell density profiles of specific cell types enables unbiased validation and refinement of known anatomical boundaries as well as the identification of novel subregions within established anatomical structures. The use of multiple well-characterized cell-type markers may facilitate future combinatorial analyses for more detailed spatial segmentation, further refining distinct areas based on unique cell type-driven brain functions.

Finally, while our study utilizes cell density analyses based on cell class- and type-specific Cre-dependent mouse models, we note that similar analyses could be conducted using whole-brain immunolabeling methods with cell type-specific antibodies. This approach may extend the utility of our method to species beyond the mouse, enabling unbiased and quantitative investigations of brain cellular architecture across evolution.

## MATERIAL AND METHODS

### Cre-reported lines

Cre driver ‘‘knock-in’’ animals were crossed with reporter mice (CAG-LoxP-STOP-LoxP-H2B- GFP) as previously described ^3^. Animal procedures were performed under the Cold Spring Harbor Laboratory Institutional Animal Care and Use Committee (IACUC) approval. All animals received food and water ad libitum and were housed under constant light and temperature conditions (12 hr cycle lights ON: 0600, lights OFF: 1800).

### Brain samples preparation and imaging of cell-type distributions

All animals were anesthetized with ketamine/xylazine and perfused transcardially with isotonic saline followed by 4% paraformaldehyde (PFA) in 0.1M phosphate buffer (PB, pH 7.4). Brains were post-fixed overnight at 4C after extraction in PFA and stored at 0.05M PB until imaging.

For imaging, brains were embedded and cross-linked with oxidized 4% agarose as previously described ^3,12,64^. Whole brain imaging of Cre animals was performed using serial two-photon tomography (STPT). The entire brain was coronally imaged at an X,Y resolution of 1µm and Z-spacing of 50 µm ^3,12,64^.

### Whole mount NeuroTrace staining

Whole brain NeuroTrace staining was performed with a modification of iDISCO+ protocol^65,66^. In order to reduce methanol induce distortions in the tissue from the original iDISCO+ protocol, samples were incubated in a solution containing 3% Triton-X in PBS with heparin for 1 week. Thereafter, samples were gently washed in PTwH 4x 8h. Finally, brain samples were incubated in a solution containing PTwH and 20% NeuroTrace for 4 weeks at 37C. Samples were finally washed in PTwH 4x 8h. Before imaging in STPT, samples were post-fixed again in 4% PFA overnight at 4C to avoid sectioning artifacts.

### Anatomical features enhancing

To improve whole brain registration, intrinsic anatomical features (IAFs) in the CCF and whole brain datasets were enhanced through preprocessing using custom MATLAB and Python scripts. First, the main IAFs in the reference brain were identified and highlighted on the reference brain, followed by the application of a Sobel operator to reduce noise and computational cost during image registration. For the imaged datasets, the stack images intensities were first homogenized to avoid signal interference during registration process using gamma compression, followed by the application of a Sobel operator to get the IAFs. This ensured that all images were consistent in format and that IAFs were enhanced identically. By enhancing IAFs in the datasets.

### Image registration

3D whole brain datasets were registered to the CCF reference brain using a three-step process. First, images were preprocessed as previously described (see *Anatomical features enhancing* above for description). Second, a 3D affine transformation was calculated using the Elastix registration toolbox^67^ with Advanced Mattes mutual information metric. This transformation was then followed by a 3D B-spline transformation to further improve the registration alignment. Accuracy was checked both visually and using brain unique identified points as previously described^1^.

### High resolution image registration transformation

After image registration at low resolution, the output transformations were used to generate high-resolution registered datasets using custom MATLAB scripts. After registration it was generated a 3D displacement field that identifying the transformation of each pixel from the original image coordinate space onto the CCF reference space. Then, the displacement field of the initial registration was transformed onto the original high resolution coordinate system and used to compute the high-resolution registration transformations from the original images.

### STPT cell counting

Automatic cell counting was performed using a convolutional neural network (CNN) trained on H2B-GFP nuclear signaling data ^3,64^. First, an unsupervised detection algorithm based on structure tensor and connected components analyses was developed to identify and segment individual cells. The results were used to generate a dataset of 270 random segmented image tiles from three different datasets, containing approximately 1350 cells from multiple brain regions.

This subset of data, both from raw and segmented with unsupervised algorithm, was then used to train the CNN for cellular detection.

Accuracy of the detection network was assessed using different datasets with sample images from multiple brain regions to account for different signal to noise ratios based on intrinsic autofluorescence. Cell detection on these datasets was done manually by 3 different experts and final accuracy calculated based on the mean values comparing the resulting output from the trained network with the expert’s manual detection.

### Hierarchical Cluster analysis

To highlight the most relevant regions for each cell type, we applied maximum normalization based on their area densities, i.e. we normalized each cell type by the maximum brain region density. We then performed a hierarchical cluster analysis with brain region correlation based on cell types distribution as pairwise distance, and ward metric to calculate the distances. All analyses were performed using custom Python scripts.

### DBSCAN cluster analysis

DBSCAN cell type density-based analysis was performed using custom Python scripts. First, we calculated, for each individual dataset, all pairs of points within a radius of 150 μm. We then calculated the 10^th^, 20^th^, 30^th^, 49^th^, 50^th^, 60^th^, 70^th^, 80^th^ and 90^th^ percentiles for each individual dataset. For each cell type we finally calculated then mean value for each percentile and then multiply by the number of datasets, obtaining the estimation of the percentiles for each cell type with all datasets. We then used the estimated percentile values to perform the DBSCAN algorithm at each percentile.

## Supporting information

Cortical density distribution

Cre driver lines labeling validation

## Acknowledgments

This study was supported by N.I.H 5U01MH114824-04 to P.O, NIH/BRAIN Initiative RF1MH128969 and UM1NS132358 to ZW, and R01NS133744 to HWD.

## Authors contributions

P.O. supervised this study. P.O. and R.M.C. designed the study, led the conceptualization, and coordinated the data generation and analysis. P.O. and J.A.H. secured funding for the research project. H.D., J.A.H., and Z.W. contributed to the conceptualization. R.M.C. conceptualized and developed the computational methods with help from J.P. and contributions from K.U.V. and C.E. Analyses were performed by R.M.C. with help from R.P. sand J.P. Animal work was performed by R.P., R.D., N.C., and K.E.H. R.P. and R.D. conducted the image setup and acquisition, under the supervision of P.O. and R.M.C. P.O. and R.M.C. wrote the original manuscript draft. All authors contributed to the review and editing of the manuscript.

---

**Supplementary Figure 1:**
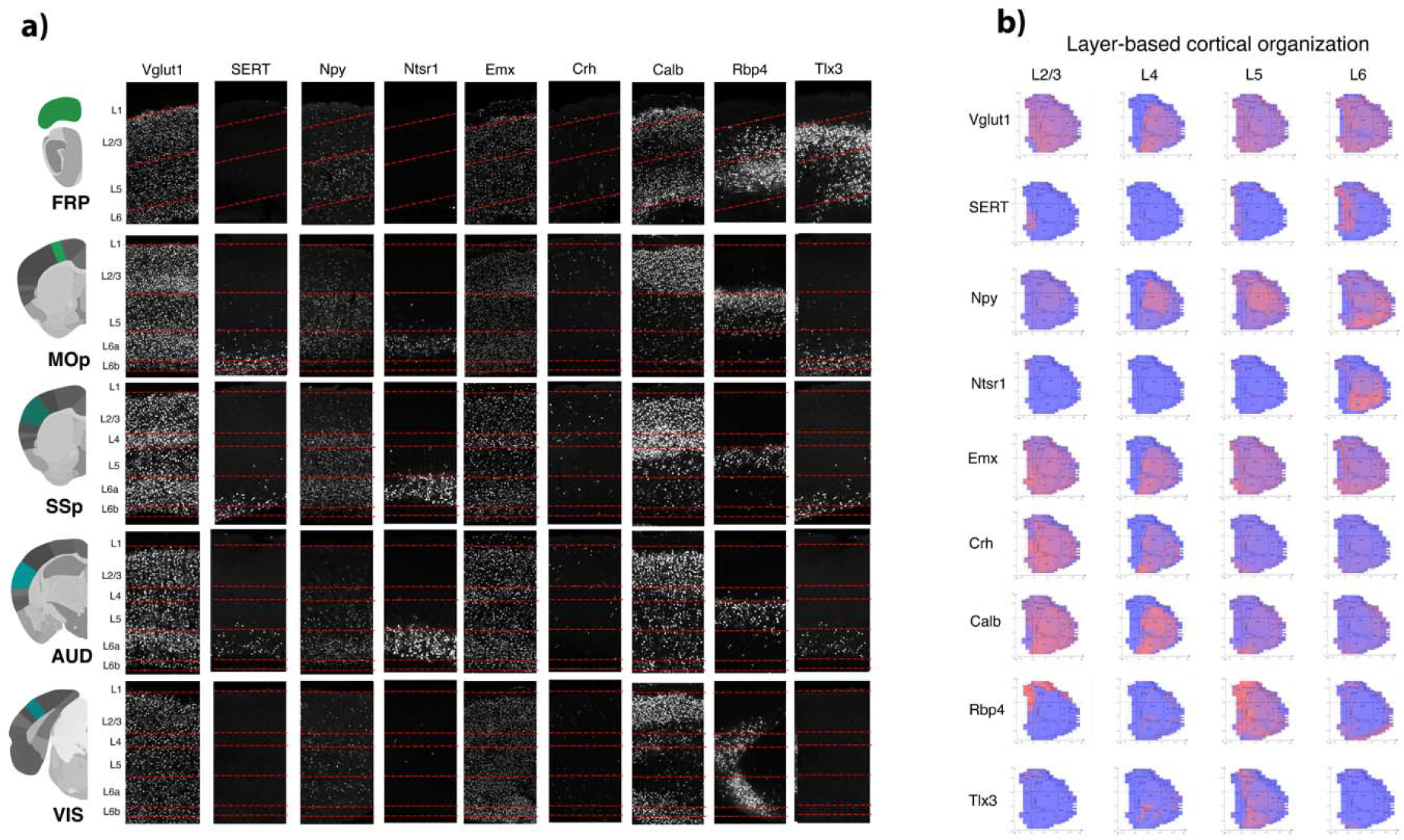
Cortical density distribution: **a)** STPT coronal sections showing the laminar distribution of nine cell type markers (VGlut1, SERT, Npy, Ntsr1, Emx, Crh, Calb, Rbp4, and Tlx3) across five cortical regions: frontal pole (FRP), primary motor cortex (MOp), primary somatosensory cortex (SSp), auditory cortex (AUD), and visual cortex (VIS). Red horizontal lines indicate cortical layer boundaries. **b)** Layer-based cortical flatmaps displaying the spatial distribution of the same nine cell types across four cortical layers (L2/3, L4, L5, L6). These flatmaps reveal both layer-specific and region-specific distribution patterns of each cell type marker.

