## Supplementary material for "A Comprehensive Atlas of Cell Type Density Patterns and Their Role in Brain Organization": Cortical density distribution

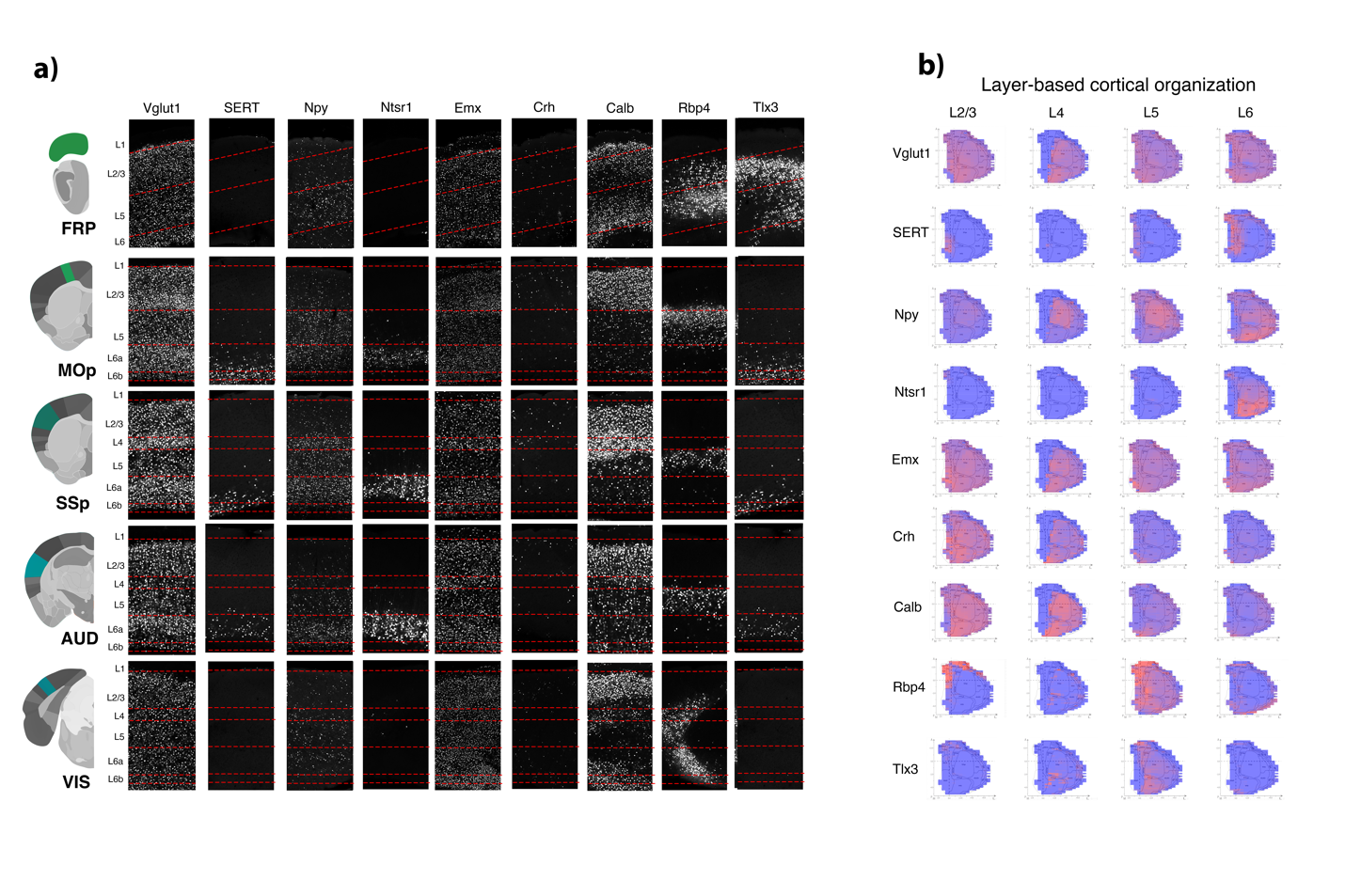

**Supplementary Figure2: Cortical density distribution: a)** STPT coronal sections showing the laminar distribution of nine cell type markers (VGlut1, SERT, Npy, Ntsr1, Emx, Crh, Calb, Rbp4, and Tlx3) across five cortical regions: frontal pole (FRP), primary motor cortex (MOp), primary somatosensory cortex (SSp), auditory cortex (AUD), and visual cortex (VIS). Red horizontal lines indicate cortical layer boundaries. **b)** Layer-based cortical flatmaps displaying the spatial distribution of the same nine cell types across four cortical layers (L2/3, L4, L5, L6). These flatmaps reveal both layer-specific and region-specific distribution patterns of each cell type marker.
