## Supplementary material for "A Comprehensive Atlas of Cell Type Density Patterns and Their Role in Brain Organization": Cre driver lines labeling validation

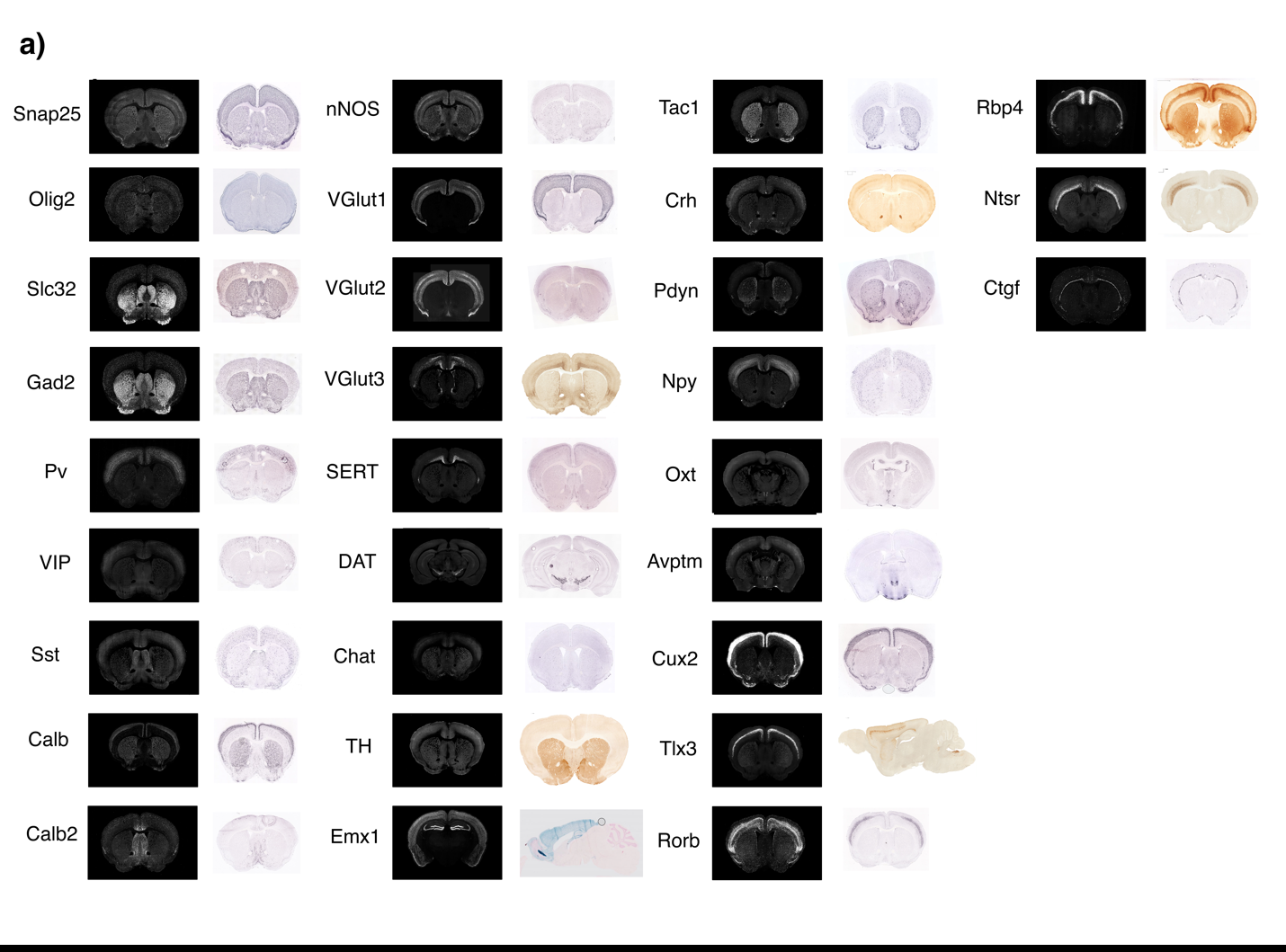

**Supplementary Figure1: Cre driver lines labeling validation.** STPT coronal planes (left panels) from the thirty Cre driver mice used in this study comparing the labeling to Allen Brain Atlas (ABA) *in situ* or GENSTAT data.
